# Acute Cytomegalovirus Infection Modulates the Intestinal Microbiota and Targets Intestinal Epithelial Cells

**DOI:** 10.1101/2022.03.17.484735

**Authors:** Vu Thuy Khanh Le-Trilling, Jana-Fabienne Ebel, Franziska Baier, Kerstin Wohlgemuth, Kai Robin Pfeifer, Aart Mookhoek, Philippe Krebs, Madita Determann, Benjamin Katschinski, Alexandra Adamczyk, Erik Lange, Robert Klopfleisch, Christian M. Lange, Viktoriya Sokolova, Mirko Trilling, Astrid M. Westendorf

## Abstract

Primary and recurrent cytomegalovirus (CMV) infections frequently cause CMV colitis in immunocompromised as well as inflammatory bowel disease (IBD) patients. Additionally, colitis occasionally occurs upon primary CMV infection in patients who are apparently immunocompetent. In both cases, the underlying pathophysiologic mechanisms are largely elusive - in part due to the lack of adequate access to specimens. We employed the mouse cytomegalovirus (MCMV) model to probe into the association between CMV and colitis. During acute primary MCMV infection of immunocompetent mice, the gut microbial composition was affected within the first 5 days post-infection as manifested by an altered ratio of the *Firmicutes* to *Bacteroidetes* phyla. Interestingly, these microbial changes incited with high-titer MCMV replication in the colon, mild crypt necrosis and increased colonic pro-inflammatory cytokine levels. Further analyses revealed that murine and human intestinal epithelial cell line as well as primary intestinal crypt cells/ organoids represent direct targets of CMV accompanied by increased cell mortality upon infection. Accordingly, *in vivo* MCMV infection disrupted the intestinal epithelial barrier and increased apoptosis of intestinal epithelial cells. In summary, our data show that CMV induces colitis in immunocompetent hosts by altering the intestinal homeostasis.

## Introduction

Cytomegaloviruses (CMVs) are prototypical members of β-*herpesvirinae*. The majority of the global adult population is latently infected with human CMV (HCMV). Although most HCMV infections in immunocompetent adults progress subclinically, fatal infections sporadically occur in apparently healthy individuals [1]. CMV colitis emerges most commonly in immunocompromised hosts due to primary infections or reactivation events of latent CMV. However, CMV colitis can also occur in healthy patients without immunodeficiency – usually in a setting of primary infection [2, 3]. The typical clinical presentation of CMV colitis includes abdominal pain, fatigue, fever, non-bloody or bloody diarrhea, and weight loss. Intriguingly, all these symptoms overlap with the clinical presentation of inflammatory bowel disease (IBD) [4, 5]. IBD constitutes a group of intestinal disorders that cause chronic inflammation of the digestive tract. The precise etiology of IBD is unknown. The prevailing hypothesis suggests that IBD results from an exaggerated immune response that is triggered by environmental factors towards altered gut microbiota or pathogenic microorganisms in a genetically prone host [6–8]. Interestingly, the prevalence of active CMV infection in the colon is considerably higher in patients with IBD compared to control populations [9] and CMV infections are associated with more severe manifestations of IBD [10, 11].

The gut microbiome exhibits many critical roles in maintaining human health [12] and is involved in the development and maintenance of the host immune system [13]. It is generally accepted that IBD is associated with microbial alterations; however, it is unclear whether such alterations are causes of the intestinal inflammation or rather represent a consequence of it. Alterations of the gut flora are also well documented in different viral infections [14, 15]. In human immunodeficiency virus (HIV) and simian immunodeficiency virus (SIV) infections, correlations between microbial dysbiosis and inflammatory cytokine production in the gut were observed and, importantly, linked to chronic immune activation [16]. For persistent CMV infection, alterations in the microbiota have been also reported. Santos Rocha *et al*. [17] found that RhCMV infections of rhesus macaques are associated with a significantly altered composition of gut microbiota and increased host immune cell activation. However, it remained unclear whether this is a long-term consequence of the conditions induced by persistent infection or whether acute CMV infection may be sufficient to trigger alterations in the gut microbiota and thereby influence the host immunity.

In the present study, we provide evidence that acute primary MCMV infection of immunocompetent mice alters the gut microbial composition and is accompanied by high-titer CMV replication in the colon, mild pathological changes in gut architecture, higher levels of colonic pro-inflammatory cytokines, and a leaky intestinal epithelial barrier. These *in vivo* findings were corroborated by studies employing murine and human primary intestinal epithelial cells in complex organoids, indicating that epithelial cells exhibit high rates of cell mortality upon CMV infection. Furthermore, immunohistochemistry of colonic tissue sections from IBD patients with a concurrent CMV replication showed that HCMV infects intestinal epithelial cells, among others. Our findings suggest a pathomechanism by which CMV infections may interfere with intestinal homeostasis und foster inflammation in the gastrointestinal tract.

## Results

### MCMV infection alters the microbial composition in the feces

While the modulation of the commensal microbiota by viruses has been described in the literature, there is a lack of information regarding the effect of primary CMV infection on the gut microbiota and associated host immunity. To investigate the consequences of an acute primary CMV infection on the microbial composition in an immunocompetent host, we infected BALB/c mice with MCMV, and collected feces samples before infection and at day 2 and 5 post-infection. DNA was extracted and the V3-V4 regions of the 16S rRNA gene were sequenced. A total of 5 bacterial phyla were detected in all analyzed samples. The predominant phyla were *Firmicutes* and *Bacteroidetes*, followed by *Proteobacteria*, *Deferribacteres* and *Actinobacteria*, which were less abundant (Figure 1 A). No significant differences in the relative abundance of *Proteobacteria*, *Deferribacteres*, *Actinobacteria* were observed between fecal samples from mice before infection (‘non-infected’) and from the same mice 2 and 5 days post MCMV infection. In contrast, at 2 and 5 days post MCMV infection, the abundance of the phylum *Firmicutes* was decreased and the phylum *Bacteroidetes* was substantially more abundant (Figure 1 A). The *ratio of Firmicutes to Bacteroidetes* has been used to express the degree of dysbiosis in the colon [18]. Of note, this ratio was reduced in infected mice, regardless of the days post-infection (Figure 1 B). When we determined whether the microbial changes in the phyla are based on alterations on family level, no changes in one specific family was detected in the *Firmicutes* phylum, rather, we observed a slight decrease in all families (data not shown). In contrast, there were higher abundances of *Rikenellaceae* and *Prevotellaceae* families in the *Bacteroidetes* phylum at day 5 post-infection compared to the mice before infection (Figure 1 C). To analyze if the observed CMV-driven dysbiosis is a long-lasting or rather a short-term effect, we infected BALB/c mice with MCMV, and collected feces samples before infection and at day 2, 7, 10 and 12 post-infection. DNA was extracted and relative abundance of the phyla *Firmicutes* and *Bacteroidetes* was analyzed via quantitative PCR. In accordance with aforementioned results, the ratio of *Firmicutes* to *Bacteroidetes* was reduced early post-infection, but this effect was completely receded at day 7 to 12 post-infection (Figure 1D), suggesting gut dysbiosis as an early transient event in MCMV infection.

**Fig. 1.**
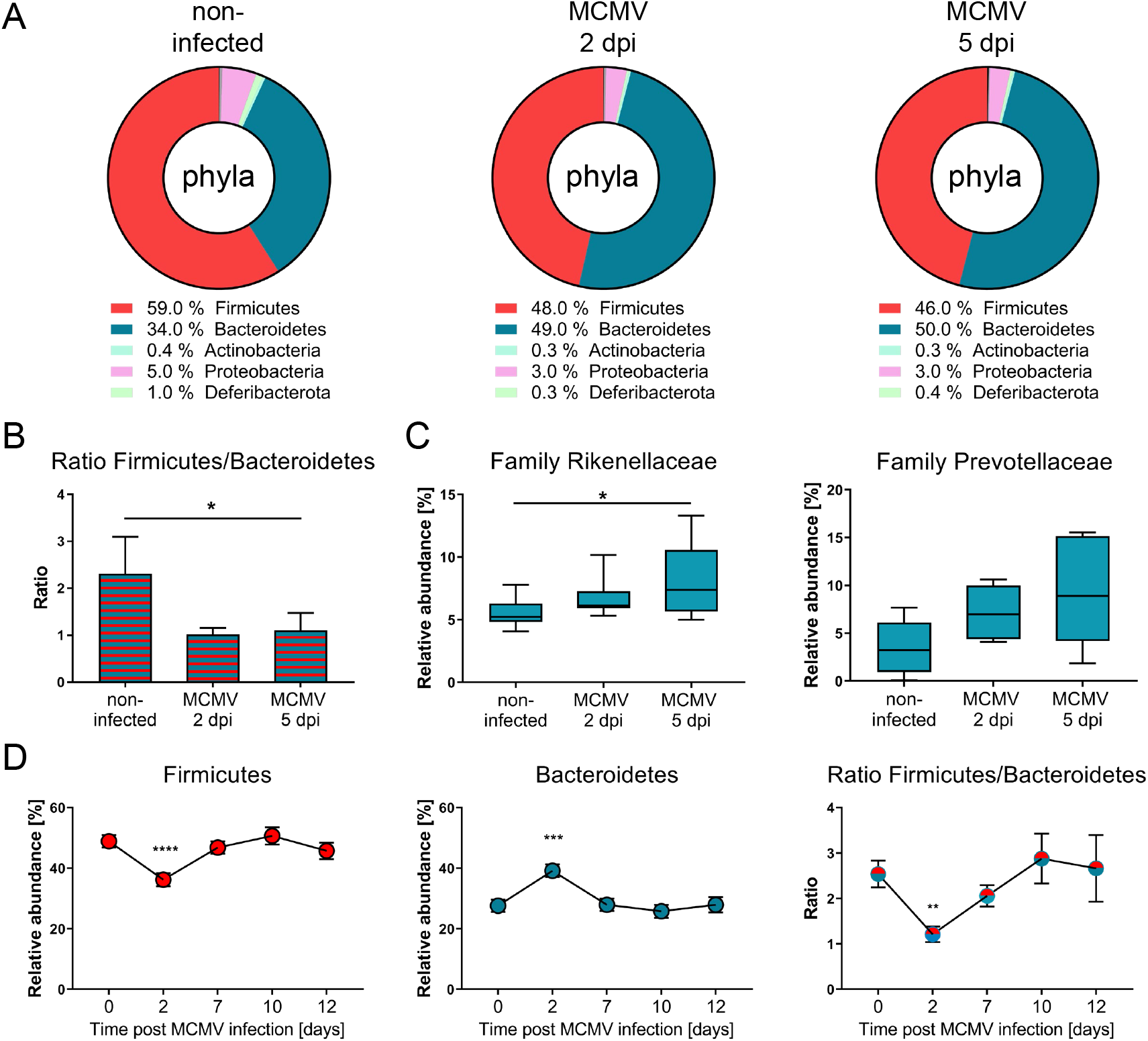
Acute MCMV infection alters the microbial composition. BALB/c mice were infected with MCMV, and feces samples were taken before infection and at indicated time points post-infection. or a quantitative PCR was performed for the phyla *Firmicutes* and *Bacteroidetes*. (A) DNA was extracted and V3-V4 regions of the 16S rRNA gene were sequenced. OTU-based profiles were analyzed using Rhea Script for R. Data was examined regarding relative abundances of the occurring bacterial phyla. (B) Ratio abundance of *Firmicutes* to *Bacteroidetes*. (C) Relative abundance of *Rikenellaceae* and *Prevotellaceae* families. (D) DNA was extracted and a quantitative PCR was performed for the phyla *Firmicutes* and *Bacteroidetes*. Relative abundance of *Firmicutes* and *Bacteroidetes* in the feces of noninfected and 2 to 12 days infected animals and Ratio abundance of *Firmicutes* to *Bacteroidetes*. Data are shown as mean ± SEM. Statistical analyses were performed using Friedman test followed by uncorrected Dunn’s multiple comparisons test or Student’s t-test. *, p < 0.05

### MCMV infection induces mild colonic inflammation with a disturbance of the intestinal barrier

A reduced ratio of *Firmicutes* to *Bacteroidetes* and an enhanced abundance of *Rikenellaceae* and *Prevotellaceae* were described to be associated with intestinal inflammation in patients suffering from IBD [18–20]. To evaluate potential direct effects of MCMV on the gut homeostasis, we first determined viral loads in different organs at day 5 post-infection. As expected from previous work [21], we observed high viral titers in the spleen, liver, and salivary glands of MCMV-infected animals. Given that the intestinal organs are not routinely assessed by most MCMV researchers, we were surprised to find that MCMV replication in the colon reached virus titers that were comparable to the amounts of virus in livers and spleens. In these different organs, MCMV titers of approximately 10^3^ - 10^5^PFU per gram of tissue were found (Figure 2 A). To determine a potential correlation between the relative abundance of different bacterila taxa and MCMV titers, we performed a Pearson correlation analysis. Correlations are plotted as colored circles representing the correlation coefficient direction (negative = red, positive = blue). The p-values are depicted by the circles’ size. Our data indicate a positive correlation between viral titers in the colon and the abundance of *Bacteroidetes* and a negative correlation with *Firmicutes* at 5 days post infection (Figure 2 B).

**Fig. 2.**
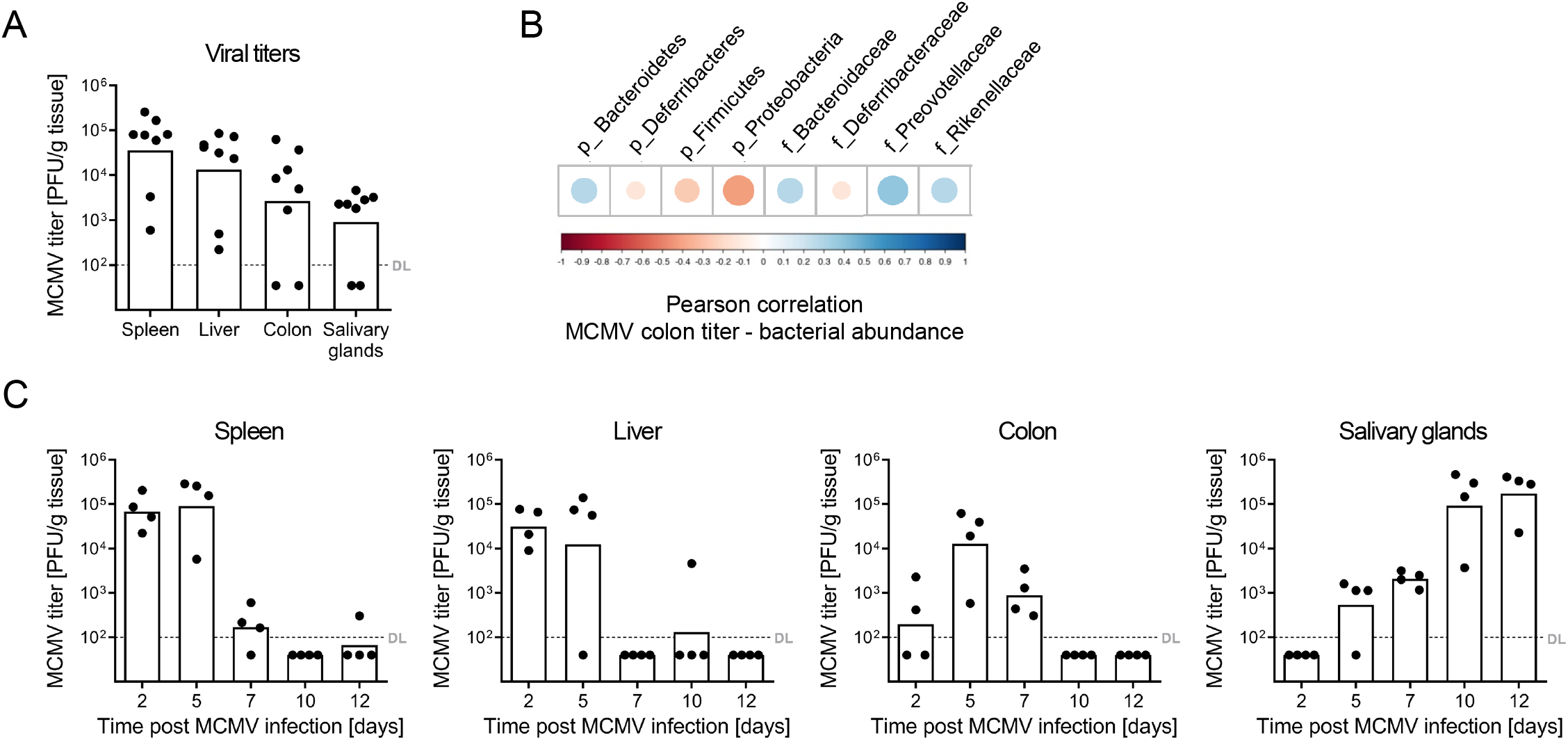
MCMV replication in the colon. BALB/c mice were infected with MCMV. (A) Viral loads in indicated organs at day 5 post-infection. (B) Pearson correlations between relative abundance of taxa and MCMV titers in the colon were calculated. Correlations are plotted as colored circles representing the correlation coefficient directions (negative in red, positive in blue). The uncorrected p-values are depicted by the circles’ size (high significance = big size). (C) Viral loads in the spleen, liver, colon, and salivary glands at indicated time points post-infection. All titrations were done in quadruplicate. Bars depict the geometric mean, dots show titers of individual mice. DL, detection limit.

To better understand the effects of MCMV in the gut, we determined the colonic viral loads in a longitudinal manner and compared the results to the viral loads in spleen, liver and salivary glands (Figure 2 C). In accordance with previous work [21], very high viral titers in the spleen and liver were detected early at day 2 and 5 post-infection with a very strong drop at day 7. In contrast, viral replication in the colon was delayed, starting at 2 days post-infection, peaking at day 5, and still profound viral titers were observed at 7 days post-infection. As expected, viral titers in the salivary glands started to increase at 5 days post-infection with increasing at later time points.

Intrigued by the profound MCMV replication in the colon, we analyzed whether the high colonic MCMV titers were associated with changes in the colonic architecture. Thus, we infected BALB/c mice with MCMV and performed histological analysis of the colon at the time point, at which viral replication was the highest (day 5 post-infection). Interestingly, mild crypt necrosis and a slightly enhanced epithelial hyperplasia were observed in MCMV-infected animals (Figure 3 A), which was associated with enhanced colonic secretion of TNF-α, IFN-γ, and in tendency enhanced IL-6 levels (Figure 3 B). Since MCMV is a cytopathic virus and IL-6 can modulate the intestinal barrier [22], we tested whether MCMV infection affects the intestinal permeability *in vivo*. Naive and 5 day MCMV-infected mice were orally gavaged with FITC-labeled dextran beads. Four hours later, the intestinal permeability was assessed by determination of FITC-dextran beads translocated from the gut lumen to the serum. Consistent with the finding that the cytopathic MCMV replicates in intestinal tissues and alters their morphology, FITC-dextran concentrations in the serum of MCMV-infected mice were significantly increased compared to naive control mice (Figure 4 A). In addition, the level of apoptosis in the colonic epithelium was determined by immunostaining of cleaved caspase-3. As shown in Figure 4 B, colon tissue sections of MCMV-infected mice exhibited a slight increase in apoptotic epithelial cells. These data provide evidence that MCMV infection affects the intestinal epithelium, causing a mild inflammation and an impairment of intestinal barrier functions in immunocompetent hosts.

**Fig. 3.**
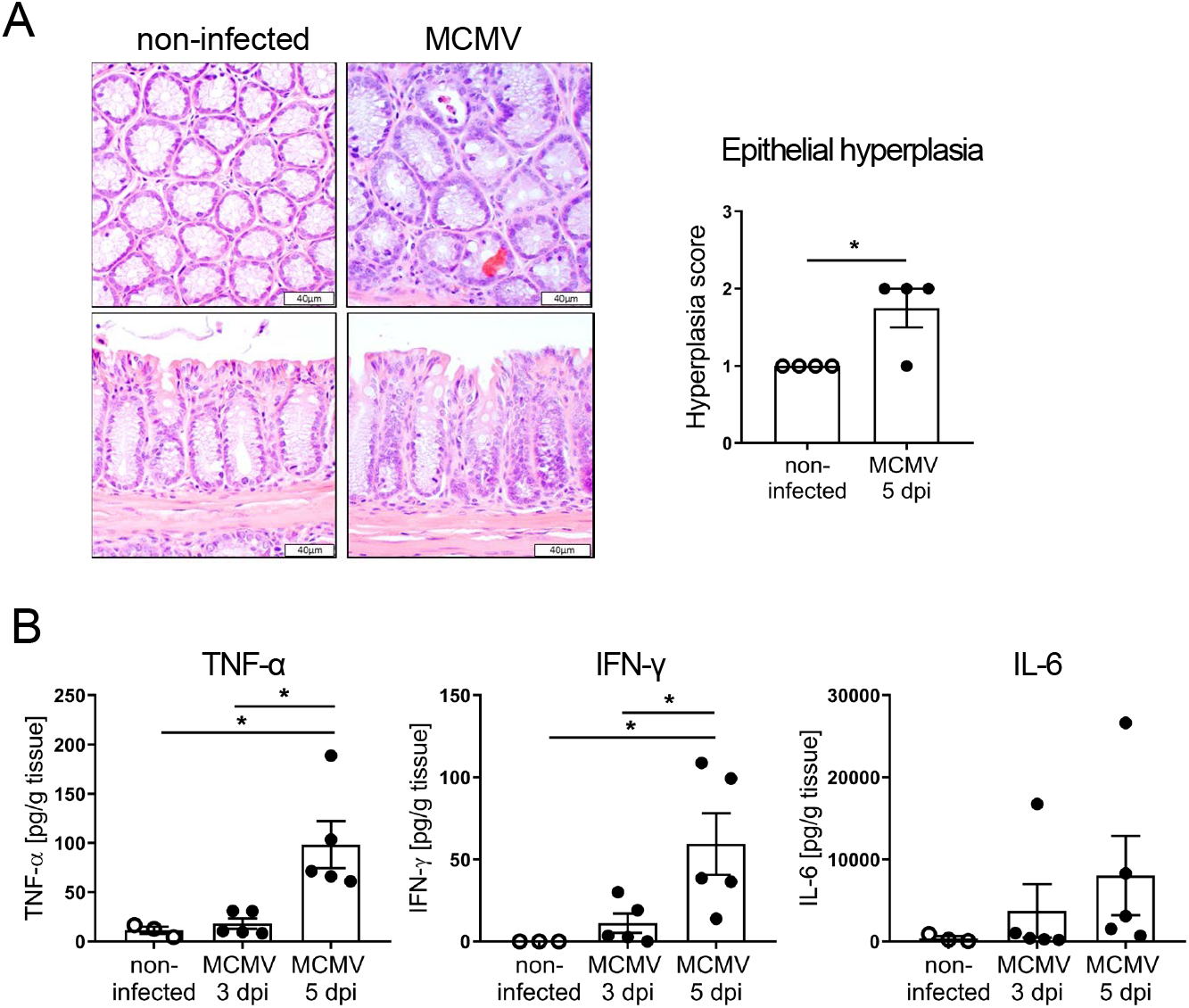
MCMV replication in the colon is associated with mild colonic inflammation. BALB/c mice were infected with MCMV. (A) Representative pictures (scale bars 40 μm) of H&E-stained colon sections of non-infected mice and mice after 5 days of infection. Epithelial hyperplasia was scored. (B) Secretion of TNFα, IFNγ, and IL-6 from *in vitro* cultured colon explants was determined by Luminex technology. Statistical analyses were performed using Mann-Whitney U test or Student’s t-test. *p < 0.05

**Fig. 4.**
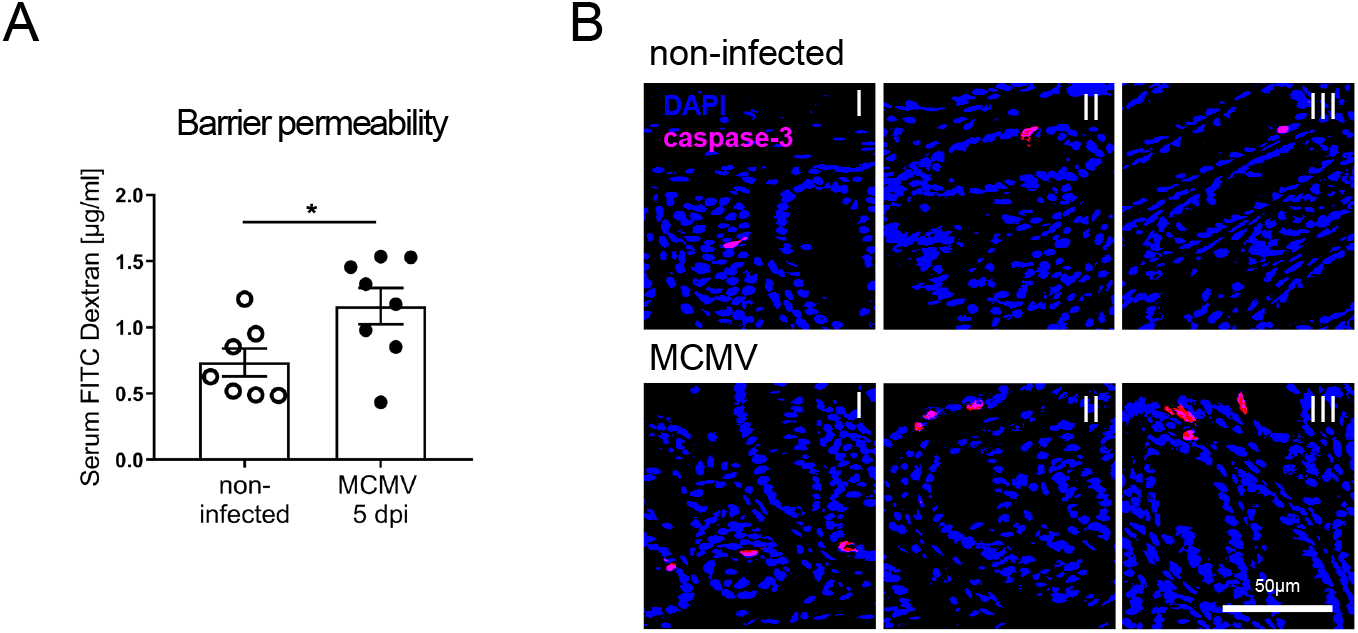
MCMV replication in the colon reduces the intestinal barrier. BALB/c mice were infected with MCMV. (A) At day 5 post-infection, each group of mice received FITC-dextran beads through oral gavage. Serum FITC-dextran concentrations were measured 4 hours after gavage. (B) Immunohistochemistry staining on colonic tissue sections of MCMV infected and non-infected mice was performed using hematoxylin (nucleus and DNA) and DAB (caspase-3). Images were dearrayed via color deconvolution, pseudocolored for nucleus/DNA (blue) and caspase-3 (red) and merged, resulting in caspase-3 positive cells (purple). Scale bar 50μm. Statistical analysis was performed using Student’s t-test. *p < 0.05

### Murine intestinal epithelial cell lines and organoids are highly susceptible to MCMV infection

To define the role of intestinal epithelial cells during gastrointestinal infection, we further investigated the susceptibility of these cells to CMV infection using a murine intestinal epithelial cell line (MODE-K) and primary intestinal organoids. First, MODE-K cells were mock-treated or infected with MCMV expressing GFP at graded virus doses (0.007 - 10 PFU/cell). As infection control, we used highly MCMV-permissive immortalized mouse embryonic fibroblasts (iMEF; described in [23]). Interestingly, MODE-K cells were nearly as susceptible to MCMV infection as the highly permissive iMEFs, as indicated by the GFP signal intensity 24 h post-infection in higher dilutions of the input virus (Figure 5 A).

**Fig. 5.**
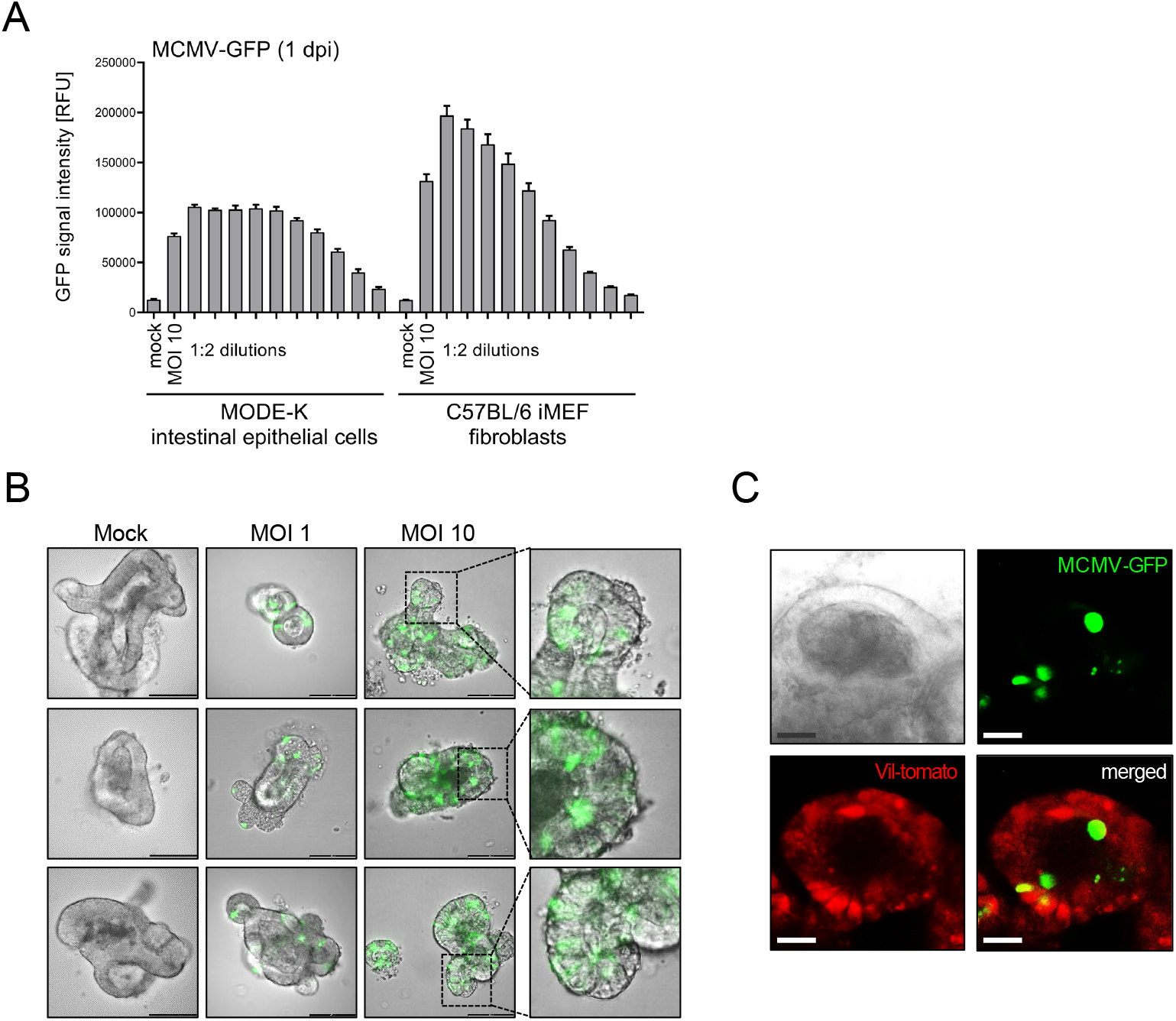
Intestinal epithelial cells are permissive for cytomegalovirus infection. (A) MODE-K cells and iMEF cells were infected *in vitro* with MCMV-GFP at indicated MOIs. Infection efficiency is shown as relative GFP signal intensity. (B) Representative images of MCMV-GFP infected intestinal organoids at 1 day post-infection. Scale 100 μm. (C) Differentiated organoids from ROSA26/LSL-tdTom x Villin-Cre (Vil-tomato), which express the tdTomato red fluorescent protein in intestinal epithelial cells, were generated. Organoids (7 days of culture) were infected with MCMV-GFP at an MOI of 10. At day 3 post-infection, immunofluorescence microscopy was performed. Scale bar 20 μm. Representative confocal images of intestinal organoids are shown. Villin-expressing epithelial cells are depicted in red. MCMV-GFP-infected cells show green fluorescence. Representative images of intestinal organoids are shown.

To bridge the gap in cellular complexity between cell culture and infections *in vivo*, we set up a protocol for the MCMV infection of intestinal organoids. Intestinal organoids are derived from self-organizing and self-renewing intestinal stem cells and closely recapitulate the native intestinal epithelium. This *ex vivo* model for the gut fully reproduces the structural architecture of the intestinal epithelium and contains all major intestinal cell lineages [24, 25]. Intestinal organoids were established by isolating intestinal crypts containing stem cells from the gut of BALB/c mice and cultured for 5-10 days until the development of differentiated organoids (Figure 5 B, Mock). Differentiated intestinal organoids were harvested and infected with MCMV-GFP at an MOI of 1 and 10. At day 2 post-infection, we identified infected cells by immunofluorescence microscopy (Figure 5 B). Different spots of infection were detected within the intestinal organoids with an enlargement of the infected host cells. To enable a visualization of epithelial cells within organoids and to infallibly ensure their intestinal origin as well as the epithelial identity, we isolated intestinal crypts from ROSA26/LSL-tdTom x Villin-Cre (Vil-tomato) mice that specifically express the red fluorescent protein tdTomato in intestinal epithelial cells (Figure 5 C). Again, differentiated intestinal organoids were harvested and infected with MCMV-GFP. At day 3 post-infection, we identified infected GFP-positive cells within the red epithelial cell layer by immunofluorescence microscopy (Figure 5 C). Taken together, our data showed that murine primary intestinal epithelial cells as single cells and in complex organoids are permissive for CMV infection.

### Survival of intestinal organoids is affected by MCMV infection

Regulated cell death programs are essential for the elimination of damaged or unwanted cells of multicellular organisms. In addition, programmed cell death has the capacity to act as defense mechanism against invading pathogens [26]. Consequently, CMV encodes multiple death inhibitors that are required for efficient viral replication [27]. To investigate how intestinal organoids react to MCMV infection in terms of organoid morphology and survival, we isolated intestinal crypts and differentiated intestinal organoids for 7 days, and infected them with increasing doses of MCMV. As control, we treated organoids with UV-inactivated MCMV at a calculated MOI of 10. In the mock treated group, 80% of the organoids were intact and showed crypt formation (‘budding’) at day 6 post-infection (Figure 6 A). In contrast, with increasing virus dose, the organoids started to die resulting in a substantial decrease in the proportion of crypt-forming organoids (Figure 6 A and B). Of note, treatment with UV-inactivated MCMV did not significantly alter the organoid morphology or induced cell death (Figure 6 A). Thus, MCMV is not able to prevent the cell death of infected intestinal organoids.

**Fig. 6.**
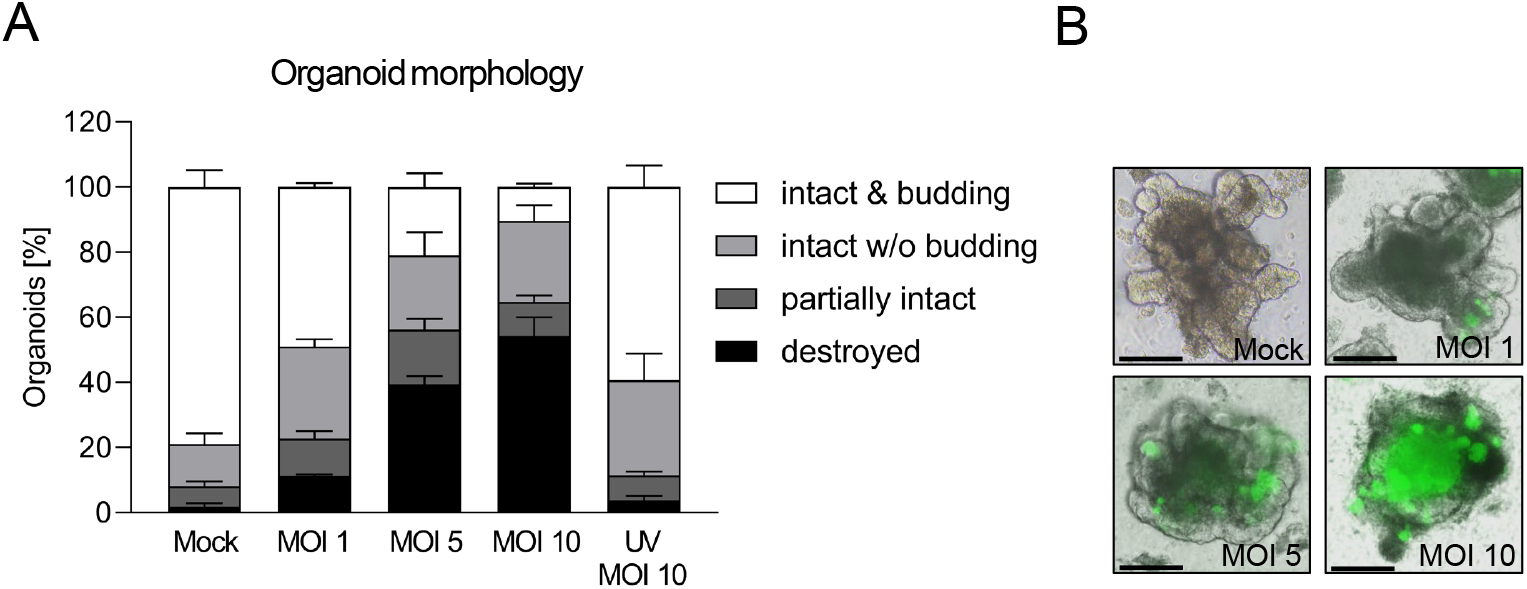
Cytomegalovirus infection impairs organoid morphology and survival. Intestinal crypts were isolated from BALB/c mice and differentiated to intestinal organoids for 7 days. Organoids were mock treated or infected with increasing MOIs of MCMV-GPF or UV-inactivated MCMV. (A) Organoid morphology was characterized at 6 days post MCMV-GFP infection. Data are shown as mean ± SEM. (B) Representative images of mock treated or MCMV infected organoids at 3 days post-infection are shown. Scale bar 100 μm.

To determine the consequences of MCMV-associated crypt necrosis and the survival of crypt stem cells *in vivo*, we isolated colonic crypts from non-infected and MCMV-infected BALB/c mice and plated these for the formation of organoids. The absolute number of colonic crypts was significantly lower when isolated from MCMV-infected mice compared with non-infected mice (Figure 7 A). The enteroid forming efficiency was determined by counting the exact number of crypts per well after plating and the number of intact organoids formed from the crypts after 24 h. Importantly, the enteroid forming efficiency was significantly reduced after MCMV infection (Figure 7 B), but the overall survival and morphology of finally formed organoids was not altered (Figure 7 C/D). Altogether, these results suggest that MCMV infection affects the survival of crypt stem cells *in vivo*.

**Fig. 7.**
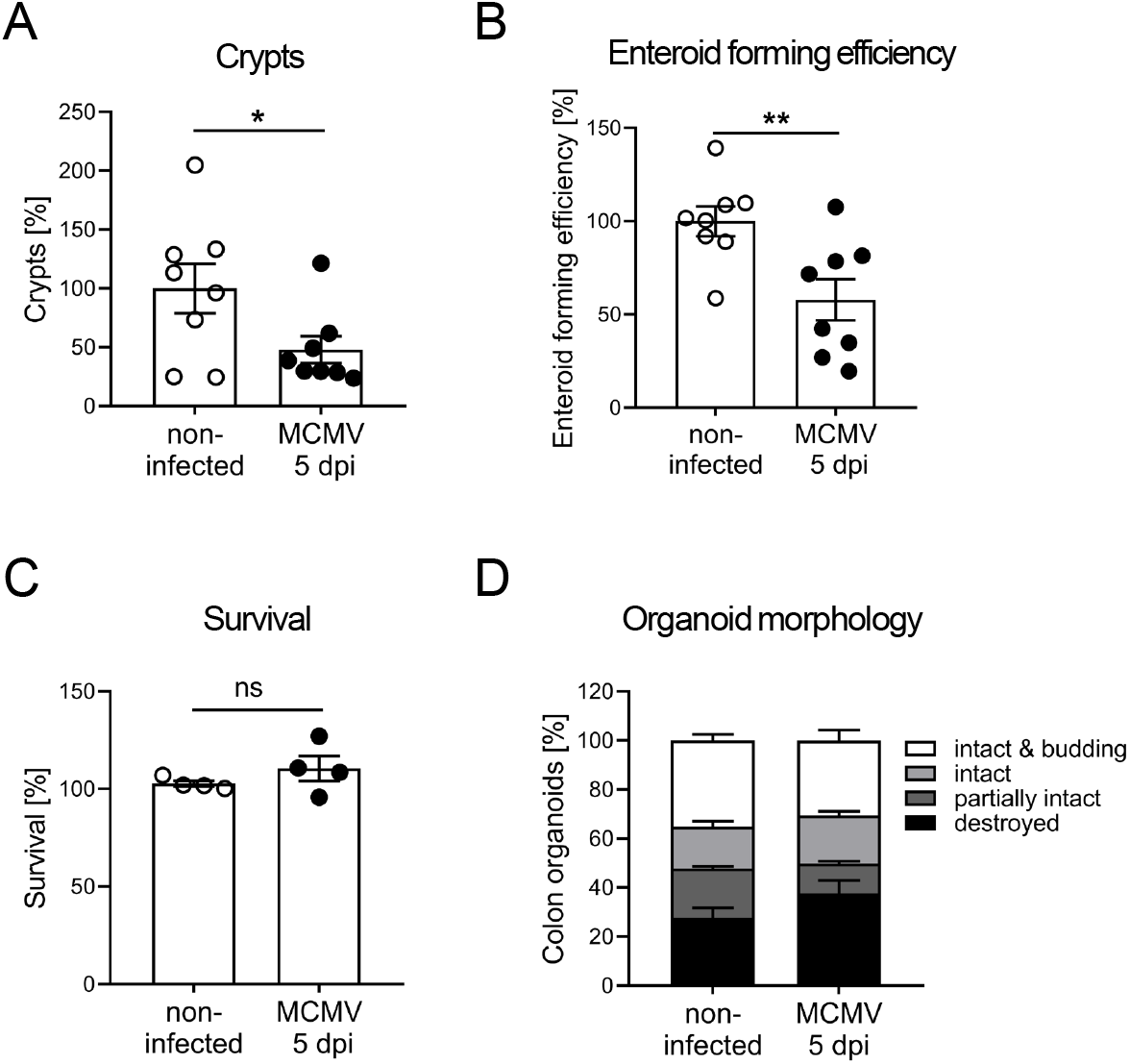
Cytomegalovirus infection *in vivo* reduces the *ex vivo* enteroid forming efficiency. BALB/c mice were infected with MCMV and at 5 days post-infection colonic crypts were isolated. (A) Relative numbers of isolated colonic crypts (the mean of the absolute number of isolated crypts from noninfected mice was set as 100%,), (B) relative enteroid forming efficiency (the mean of the relative enteroid forming efficiency of non-infected mice was set as 100%), (C) relative organoid survival and the (D) organoid morphology were determined. Data are shown as mean ± SEM. Statistical analyses were performed using Student’s t-test. *p < 0.05; **p < 0.01

### Human intestinal epithelial cell lines and organoids are highly susceptible to HCMV infection

To address the general relevance of our findings from murine studies, we assessed whether primary human intestinal epithelial cells are permissive for HCMV infection by using human gut-derived organoids. Upon full differentiation, organoids were infected with HCMV expressing GFP and at 3 days post HCMV exposure, infection was analyzed by immunofluorescence microscopy. Similar to the results with murine organoids, different spots of infection were detected within the intestinal organoid with an enlargement of the infected host cells and first signs of cell death (Figure 8 A).

**Fig. 8.**
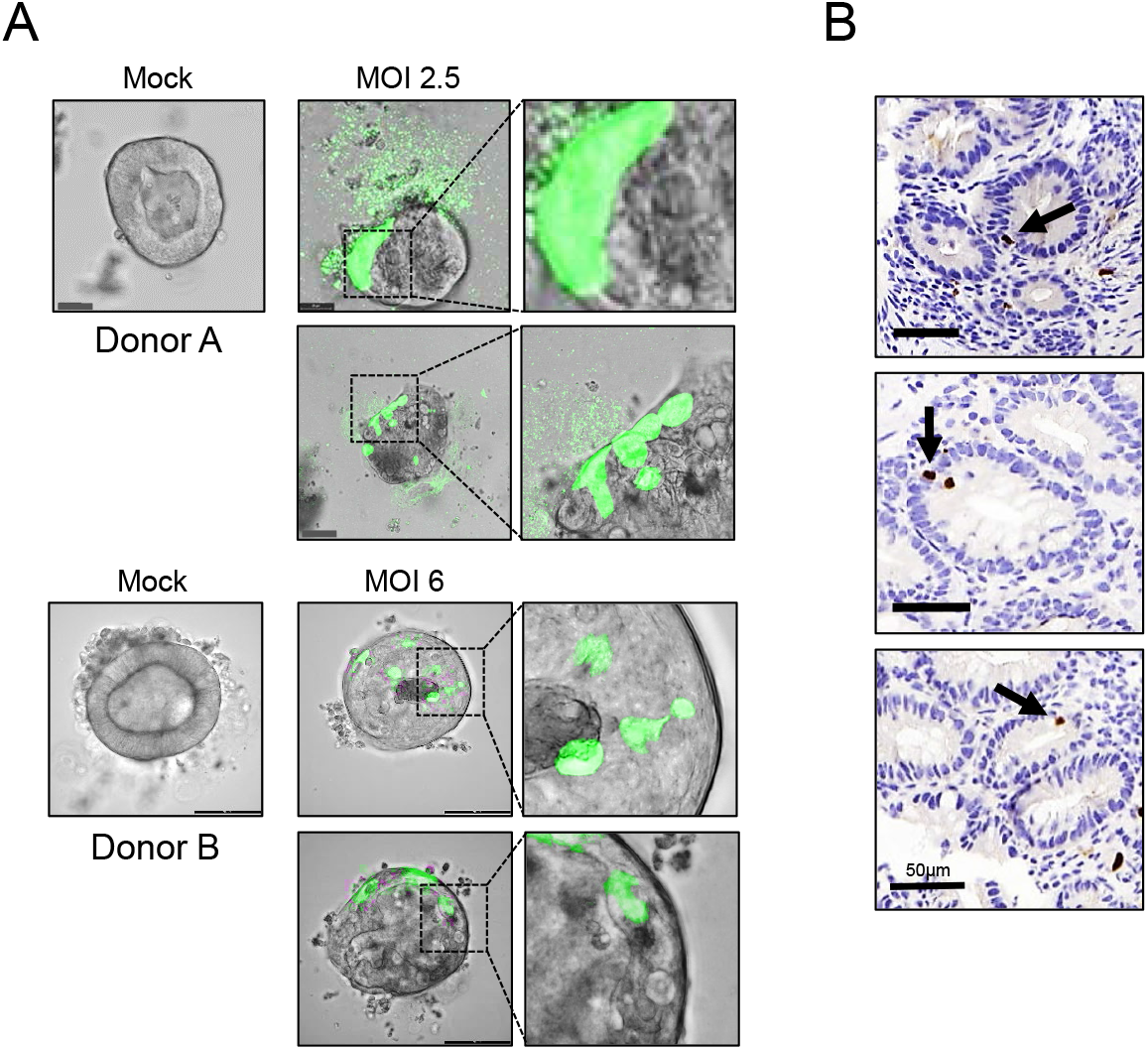
Human intestinal epithelial cells are permissive for HCMV infection. (A) Differentiated human organoids were generated from colon biopsies of two different healthy donors, mock treated or infected with HCMV-GFP at an MOI of 2.5 (donor A) or MOI of 6 (donor B). At day 3 post-infection, immunofluorescence microscopy was performed. Representative images of intestinal organoids are shown. Scale bar 20 μm for donor A, 100 μm for donor B. (B) Colonic tissue sections from IBD patients with CMV association were stained with mouse anti-human CMV. Scale bar 50μm.

Next, we performed immunohistochemistry staining for HCMV in colonic sections to determine the colonic target cells for CMV. To increase the number of colonic spots of infection we used sections from IBD patients with a concurrent CMV replication. Consistent with our *in vitro* studies, we observed, although in rare cases, the presence of HCMV-positive intestinal epithelial cells (Figure 8 B). In summary, our findings indicate that murine and human primary intestinal epithelial cells are permissive for CMV entry and gene expression with an increased cell mortality upon CMV infection.

## Discussion

CMV infections are usually controlled and confined by the adult immune system of the healthy host. However, CMV can invade end organs and can cause tissue-invasive diseases such as colitis, gastritis, or enteritis [28]. CMV diseases mainly occur in immunocompromised patients, but cases have also been reported in apparently immunocompetent patients, especially after primary CMV infection [2, 3]. The gastrointestinal mucosa is a major site of opportunistic CMV replication, causing or exacerbating intestinal inflammation that may result in severe end-organ dysfunction [29, 30]. Although the clinical manifestations of intestinal CMV infections are well described, the pathophysiologic mechanisms by which CMV promotes intestinal inflammation and the role of the microbiome in this process remain poorly understood. Especially in light of the clinical relevance of CMV colitis, it is surprising that MCMV replication in the intestine has been largely neglected. Accordingly, the gastrointestinal tract is not mentioned as a potential organ for MCMV replication in the standard reference protocol collection for MCMV *in vivo* analyses [31]. Sterility problems that may arise in cell culture caused by the gut microbiome may be one reason for this.

Even though the relative contribution may vary among different viruses, the influence of the microbiota on the susceptibility to viral infections and the outcome of infectious diseases is undisputable. Santos Rocha *et al*. [17] analyzed the microbiota of RhCMV-infected rhesus macaques. Interestingly, they identified significantly altered gut microbiota and increased host immune cell activation in subclinically infected animals. However, they could not differentiate whether this is a long-term effect of persistent infection or due to an acute CMV infection. Here, we showed that an acute primary CMV infection of immunocompetent mice is sufficient to induce alterations in the microbial composition. In more detail, the relative abundance of the phylum *Firmicutes* was reduced and the abundance of bacteria belonging to the phylum *Bacteroidetes* was enhanced. The fecal ratio of *Firmicutes* to *Bacteroidetes* has been used as proxy for the health status of a host [32]. Interestingly, we found that the ratio of *Firmicutes* to *Bacteroidetes* was significantly reduced during acute CMV infection at day 2 and 5 postinfection but completely resolved from day 7 post-infection. Recent evidence indicates that such dysbiosis is associated with the appearance of a number of inflammatory disorders and systemic diseases including gastrointestinal diseases such as IBD [18–20, 33]. Thus, acute primary CMV infection may temporarely alter the microbial community towards a composition that fosters intestinal inflammations. In line with this hypothesis, we identified high viral titers in the colon of MCMV-infected immunocompetent mice, associated with mild crypt necrosis and slightly enhanced hyperplasia of the intestinal epithelial cells during acute primary MCMV infection. In this context, CMV infection is of particular interest with regard to IBD as the prevalence of active CMV infection in the colon is considerably higher in patients with IBD relative to control patients [9] and CMV infection seems to be associated with more severe IBD [10]. Similarly, we observed HCMV positive intestinal epithelial cells in an IBD patient associated with HCMV, but the existence of a correlation/association does not necessarily inidcate causality. Nevertheless, CMV infection experiments in mouse models of IBD have shown that acute and latent CMV infection exacerbates intestinal inflammation [34–37]. Although the underlying mechanism are not entirely understood, it was demonstrated that concomitant MCMC infection increased gross bleeding [34], induced the production of pro-inflammatory cytokines or chemokine ligands in the colon [35, 37], or stimulated gut immune responses to the gut microbiota [36].

It is still of debate which cells in the colon are targeted by CMV infection. Replication of the virus in endothelial cells could generate vasculitis associated with microvessel thrombosis and local ulceration [38, 39]. Furthermore, the recruitment of CMV-infected monocytes to the mucosa that promotes the dissemination of inflammatory macrophages in the inflamed tissue was reported [40, 41]. We detected mild colonic crypt necrosis in MCMV-infected immunocompetent mice suggesting epithelial cells as a target for CMV. Necrosis is a passive but uncontrolled process, initiated by external factors such as viral infections and is characterized by a rapid breakdown of the cell membrane, resulting in the release of intracellular compounds into the extracellular space with concomitant activation of the immune system [42]. Accordingly, we measured elevated levels of colonic TNF-α, IFN-γ, and IL-6 in MCMV-infected mice. To define the role of intestinal epithelial cells during gastrointestinal infection in more detail, we made use of primary intestinal organoids. Interestingly, primary murine and human organoids are permissive for CMV infection. CMV is known to encode multiple death inhibitors that are required for efficient viral replication [27]. Nevertheless, CMV was not able to prevent the cell death of infected intestinal organoids in our experimental setup. Maidji *et al*. [43] developed a SCID-hu gut mouse model where the authors subcutaneously implanted human fetal gut into immune-deficient SCID mice. This fetal intestine developed into a differentiated human intestine with a lumen and morphologically precise mucosal layers within 4 weeks. At this stage, the authors intraluminally inoculated the human gut with CMV. Interestingly, they observed substantial CMV infection with marked mucosal damage after 7 days and a rapid depletion of epithelial cells from the fetal intestine differentiated *in vivo*. Accordingly, we could show a reduction in the *in vitro* enteroid forming efficiency in MCMV-infected immunocompetent animals compared to non-infected animals. Necrosis or programmed cell death of infected intestinal epithelial cells *in vivo* interferes with the integrity of the intestinal barrier [44]. Furthermore, IL-6, which is secreted in the colon during CMV infection, was described to decrease the intestinal barrier [22]. Consequently, it was shown that CMV infection disrupts the tight junctions of polarized epithelial cells and the adherent junctions of polarized endothelial cells *in vitro* [43]. We could now demonstrate that MCMV infection of immunocompetent mice significantly enhances the intestinal permeability *in vivo*, thus favoring the translocation of commensals and antigens from the lumen to the *lamina propria*, thereby boosting intestinal inflammation. In summary, our results suggest that primary infection with CMV modulates the mucosal integrity and, consequently, alters the risk for intestinal inflammation and may predispose to the development of IBD. In addition, the surprisingly efficient MCMV replication in the gut and the intestine-specific replication kinetics raises numerous intriguing future questions, e.g., regarding tissue-specific immune responses as well as viral adaptations to this special niche.

## Material and Methods

### Human colon biopsies

Colonic biopsies were provided from two healthy donors. Informed consent was obtained from all donors. Ethical approval was provided by the Medical Faculty of the University of Duisburg-Essen (AZ 21-9864-BO). Stained tissue sections from IBD patients with CMV association were provided by the Tissue Bank Bern (Switzerland), and images were selected among 20 anonymized cases.

### Mice

All mice were 8 to 15 weeks old, bred and housed in accordance to the guidelines of the Laboratory Animal Facility of the University Hospital Essen. Animal experiments were performed in accordance to the ethical principles and federal guidelines and approved by the Landesamt für Natur, Umwelt und Verbraucherschutz (LANUV, Germany). BALB/c were obtained from Envigo RMS GmbH. ROSA26/LSL-tdTom were obtained from the Jackson Laboratory and crossed to Villin-Cre mice [45].

### Cells, viruses, and virus titration

Primary mouse fibroblasts (mouse embryonic fibroblasts [MEF] and mouse newborn cells [MNC]) were isolated from mouse embryos and newborns, respectively, according to described protocols [46]. Mouse fibroblasts were grown in Dulbecco’s minimal essential medium (DMEM) supplemented with 10% (v/v) FCS, 100 μg/ml streptomycin, 100 U/ml penicillin, and 2 mM glutamine. Murine MODE-K[47] cells were cultivated in Roswell Park Memorial Institute (RPMI)-1640 supplemented with 10% (v/v) FCS, 100 μg/ml streptomycin, 100 U/ml penicillin, and 2 mM glutamine. All cell culture media and supplements were obtained from Gibco (Life technologies, Darmstadt, Germany). The wt-MCMV was reconstituted from the MCMV-BAC described in Jordan *et al*. [48]. MCMV-GFP has been previously described[49, 50]. MCMV propagation and determination of viral titers by standard plaque titration were performed using primary MEF or MNC [46]. All *in vitro* infections and titrations were conducted with centrifugal enhancement (900 g for 30 min). For *in vivo* infections, mice were infected intraperitoneally with 2*10^5^ PFU MCMV per mouse. Organs of infected mice were harvested, snap frozen in liquid nitrogen, and stored at −80°C until titrations were performed. HCMV-GFP (TB40EΔgpt EGFP[51]) was propagated as described in Le-Trilling *et al*. [52]. Viral GFP reporter gene expression was quantified by use of a microplate multireader (Mithras LB 943; Berthold Technologies GmbH & Co.KG, Bad Wildbad, Germany).

### DNA extraction and sequencing

Feces samples were collected and stored at −80°C. DNA was extracted using Zymo Biomics DNA miniprep kit (Zymo, Freiburg, Germany). Genomic DNA was used for qPCR analysis [53] or prepared for sequencing following the protocol #15044223 Rev. B provided by Illumina technical support: Regions V3 and V4 of the 16S rRNA gene were amplified, and samples were labelled with index primers using Nextera XT Index kit (Illumina, San Diego, CA, USA). Paired end sequencing was performed with the Illumina MiSeq system. Sequenced raw data was re-multiplexed employing the Perl programing language. By utilizing the Integrated Microbial Next Generation Sequencing [54] platform, reads with a similarity of at least 97% were assigned to operational taxonomic units (OTUs). OTU-based microbial profiles were built as described by Lagkouvardos *et al*. [55]. The OTU-based profiles where examined regarding sequencing depth, alpha diversity, relative abundances of different OTUs assigned to taxonomic levels within each sample, beta diversity, and diversity between different groups of samples. Also, Pearson correlations between the relative abundances of taxa and the MCMV titers were calculated. Analysis were performed using the Rhea script pipeline [56] for the software R.

### Histopathological analysis

Full-length colons were stored in 4% (w/v) paraformaldehyde and embedded in paraffin. Sections (4 μm) were prepared from paraffin-embedded blocks, stained with hematoxylin and eosin (H&E) and evaluated in a blinded manner according to standard techniques as described previously[57]. Caspase-3 staining was performed as described earlier[58]. Images were dearrayed using Fiji ImageJ (Version 1.52i) and the color deconvolution plugin, resulting in individual channels for hematoxylin (H); nucleus and DNA) and 3,3′-Diaminobenzidine (DAB; caspase-3). Specific threshold was set for DNA and caspase-3 and images were pseudo-colored in blue (DNA) and red (caspase-3), respectively, after which images were merged.

Human tissue sections were fixed in 4% formaldehyde and embedded in paraffin. All staining reactions were performed by automated staining using a BOND III autostainer (Leica Biosystems, Wetzlar, Germany). For immunohistochemistry, sections were deparaffinized and antigen was retrieved using 1 mM Tris solution (pH 9.0) for 30 min at 95°C. Sections were stained with mouse anti human CMV (Dako, Agilent technologies, Glostrup, Denmark) primary antibody. Specific binding of primary antibodies was visualized using a polymer-based visualizing system with horseradish peroxidase as the enzyme and 3,3-diaminobenzidine (DAB) as a brown chromogen, (Bond^™^ Polymer Refine DAB Detection from Leica Biosystems). Finally, the samples were counterstained with haematoxylin and mounted with Eukitt (O. Kindler GmbH, Freiburg, Germany) before scanning. After staining, slides were scanned using a Pannoramic 250 digital scanner (3DHISTECH).

### Colon explant culture and cytokine detection

A small explant (15-25 μg) from the distal part of the colon was cultured for 6 h in IMDM complete medium (IMDM containing GlutaMax^™^ -I, 25 mM HEPES, 10% FCS, 100 μg/ml streptomycin, 100 U/ml penicillin, 25 μM β-mercaptoethanol). Cytokine levels in the supernatants were measured by Luminex technology (R&D Systems, Wiesbaden, Germany) on a Luminex 200 instrument using the Luminex IS software (Luminex Corporation, MV’s-Hertogenbosch, Netherlands). Cytokine concentration was normalized to the respective weight of the colon biopsies.

### FITC-dextran intestinal permeability assay

Non-infected and MCMV-infected mice were orally gavaged with 150 μl of 100 mg/ml 4 kDa FITC-dextran-labeled dextran beads (Sigma-Aldrich/Merck, Darmstadt, Germany) in PBS 4 hours prior to sacrifice. FITC-derived fluorescence was quantified in the serum using a microplate multireader (Mithras LB 943). Concentrations were determined using a standard curve generated by serial dilution of FITC-dextran.

### Crypt isolation from murine intestine

Small intestinal samples were rinsed with ice-cold PBS and cut into 5-mm pieces. Tissue segments were washed multiple times with cold PBS and incubated in 5 mM EDTA-PBS for 30 min gently shaking on ice. To release crypts, tissue pieces were shaken twice vigorously in cold PBS for 30 s. The crypt suspension was passed through a 100-μm cell strainer, pelleted at 300 g for 5 min and washed at 60 g for 3 min to remove single cells. Crypts were plated in 40 μl matrigel (Corning, Tewksbury, MA, USA) and cultured with 500 μl/well murine IntestiCult^™^ Organoid Growth Medium (OGM, Stemcell Technologies, Cologne, Germany). Medium was changed every 2-3 days. MCMV infection was performed after 7 days of culture.

### Crypt isolation from murine colon, enteroid forming efficiency and survival

Colonic crypt isolation was performed as described previously [59]. Briefly, the distal colon was flushed with ice-cold PBS containing antibiotics and minced into 2-mm pieces. Tissue fragments were digested in 10 ml DMEM containing 1% FCS, antibiotics and 500 U/ml collagenase IV. Crypts were released by shaking and pipetting the tissue fragments vigorously. Isolated crypts were passed through a 100-μm cell strainer, pelleted, washed and embedded in matrigel. After polymerization, 500 μl/well of murine IntestiCult^™^ OGM were added. The enteroid forming efficiency was determined by counting the exact number of crypts after plating and the organoids formed from the crypts after 24 h. For enteroid survival, the amount of intact organoids was determined every second day for one week after crypt isolation.

### Crypt isolation from human colon biopsies

Colonic biopsies were washed several times with ice-cold PBS and incubated in 2.5 mM EDTA-PBS at 4 °C for 40 min. Tissue fragments were suspended vigorously in ice-cold PBS with a bovin serum albumin (BSA) coated 10 ml pipette. This step was repeated three times and the supernatants containing crypts were transferred to a BSA-coated tube. Crypts were pelleted at 1600 rpm for 3 min, plated in matrigel and cultured with 500 μl/well human IntestiCult^™^ OGM (Stemcell Technologies). Medium was changed every 2-3 days and organoids were passaged once a week. After the second passage, organoids were used for HCMV infection.

### CMV infection of organoids and assessment of organoid morphology

Organoids were collected in 500 μl/well cell recovery solution (Corning) and incubated for 30 min on ice to depolymerize the matrigel. The organoids were pelleted at 300 g for 3 min and washed with DMEM twice. Afterwards, organoids were resuspended in virus standard buffer (VSB) (Mock), MCMV-GFP, MCMV-m*Cherry*, UV-irradiated MCMV-GFP or the human CMV strain TB40EΔgpt EGFP diluted in DMEM for different multiplicities of infection (MOI) and incubated for 30 min at 37°C. Thereafter, samples were briefly placed on ice, embedded in matrigel and cultured with IntestiCult^™^ Organoid Growth Medium. For morphology, organoids were classified into four groups (intact budding, intact not budding, partially intact, and destroyed organoids) and counted by light microscopy at 6 days post-infection.

### Microscopy

Fluorescence microscopy was performed on an AxioObserver Z1 widefield microscope using a 20x objective and Axiocam 506 mono camera and on Leica Thunder Imager. Images were taken with Leica THUNDER imager using a 20x and 40x objective (Leica DFC9000 GT). Images were processed using computational large volume clearing method. Confocal laser scanning microscopy (CLSM) was performed with a TCS SP8 AOBS system operated by the LASX software (Leica Microsystems). Images were acquired with HC PL Fluotar 10x/0.3, HC PL APO 20x/0.75 CS2, and HCX PL Apo 63x/1.4 oil objectives. Imaging conditions: pinhole 1 AU, laser intensity 5%. The spheroid images were all recorded with the same set of parameters.

### Statistics

Statistical analyses were performed in R or in GraphPad Prism. All statistical tests were performed with significance and confidence level of 0.05.

## Data availability

16S rRNA gene data are deposited in the NCBI SRA (SRA: BioProject: PRJNA763375)

## Acknowledgement

We kindly thank Mechthild Hemmler-Roloff for excellent technical assistance. The authors are very grateful for a very generous donation from Alantra, based on which the Leica THUNDER Imager was purchased. We thank also the Imaging Center Campus Essen (ICCE), Center of Medical Biotechnology (ZMB), and University of Duisburg-Essen for access to the confocal laser scanning microscopes.

